# *FveTRM5* plays a critical role in regulating fruit shape in woodland strawberry

**DOI:** 10.1101/2025.03.10.642363

**Authors:** Zhenzhen Zheng, Liyang Wang, Qi Gao, Shaoqiang Hu, Chunying Kang

## Abstract

Cultivated strawberry is a globally important fruit crop with high economic value. Fruit shape contributes to fruit quality and diversity and is a target for breeding, but very few regulatory genes have been reported in strawberry. Here, we identified an ethyl methanesulfonate (EMS) *round fruit* (*rf*) mutant that produces round or flat fruits in woodland strawberry. The primary candidate point mutation is located in the second exon of FvH4_2g22810, causing a premature stop codon at residue 266. This gene encodes a protein with a high similarity to TON1 RECRUITING MOTIF 5 (TRM5) and has therefore been named *FveTRM5*. Transformation of *FveTRM5*pro:*FveTRM5* into *rf* could rescue the round fruit phenotype, suggesting that *FveTRM5* is responsible for *rf*. Overexpression of *FveTRM5* produced elongated organs in both Arabidopsis and woodland strawberry, suggesting a conserved role in different species. *FveTRM5* is ubiquitously expressed with higher levels in developing organs. Observation of cell shape showed that *FveTRM5* promotes cell elongation and inhibits cell division in the medial-lateral direction in the receptacle. The FveTRM5 protein localized to microtubules. In conclusion, our results suggest that FveTRM5 plays an essential role in regulating strawberry fruit shape by influencing cell elongation and cell division, providing an excellent target gene for breeding new fruit shape cultivars.

## 1. Introduction

Fruit shape is an important external quality trait that strongly influences consumer preference. The organization of floral organs and which part can develop into fruit flesh has evolved differently among species [1, 2]. The diversity of fruit shapes arises from the meristem activity and the growth patterns along the adaxial-abaxial, proximal-distal, and mediolateral axes during ovary and fruit development [1, 3, 4]. Understanding the regulation of fruit shape is of great importance for the genetic improvement of fruit crops to create new cultivars.

Three gene families are known to play key roles in the organ shape control in different plant species, including the TONNEAU1 Recruiting Motif proteins (TRMs), the Ovate Family Proteins (OFPs), and the IQ67 domain-containing proteins (IQDs) [5-7]. Mutation or overexpression of TRMs can cause altered shapes of various organs, including seeds/grains/fruits in Arabidopsis [8, 9], rice [10, 11], wheat [12], maize [13], tomato and cucumber [14, 15]. In the OFP family, OVATE and SlOFP20 in tomato are negative regulators of fruit elongation [14, 16], and *PpOFP1* overexpression due to genomic inversion results in flat fruits in peach [17-19]. The IQD protein SUN1, a calmodulin-binding protein, promotes fruit elongation in tomato [20]. A number of other OFPs or IQDs have also been reported to influence organ shape [21].

Fruit shape is closely related to cell division and cell expansion in different directions. The TRM proteins affect both cell division and cell expansion [9, 10, 12, 14, 15]. A common mechanism underlying this biological process is the modulation of the microtubular cytoskeleton. TRMs can recruit TON1 and phosphatase 2A to the microtubule arrays and the preprophase bands to control the direction of cell division [8, 22-25]. TRMs can also interact with OFPs to antagonistically modulate microtubule arrangement [6, 14]. Tomato SlTRM5 localizes to the microtubules, whereas SlOFP20 localizes to the cytoplasm and nucleus; when *SlTRM5* and *SlOFP20* were transiently expressed together in *Nicotiana benthamiana* leaves, SlOFP20 was largely translocated to the microtubules [14]. SUN1/SlIQD12 also localizes to microtubules to alter their structure, leading to changes in cell division patterns [26]. In this process, IQDs may interact with calmodulin (CaM) to sense and respond to the calcium signals [27]. The TRM-OFP module may coordinate with IQDs on the microtubular cytoskeleton to regulate organ shape [5, 6].

Cultivated strawberry (*Fragaria*×*ananassa*, octoploid) is a globally important fruit crop with high economic value. Fruit shape contributes to fruit quality and diversity and is therefore a selected trait in strawberry breeding. Unlike most fruit crops, strawberry flesh develops from the enlarged tip of the stem below the floral organs, known as the receptacle [2]. Both wild and cultivated strawberries have different fruit shapes, ranging from flat to long conical [28, 29]. Fruit shape in strawberries responds to environmental stimuli and endogenous factors such as auxin and gibberellin (GA) levels [30, 31]. Some quantitative trait loci (QTLs) responsible for natural shape variation in strawberry have been identified [32, 33], but few regulatory genes were reported.

Woodland strawberry (*F. vesca*) is a wild diploid species that typically produces small conical fruits. In this study, we isolated the fruit shape control gene, *FveTRM5*, from the ethyl methanesulfonate (EMS) mutant *round fruit* (*rf*) in woodland strawberry. Genetic complementation and overexpression assays validated the important functions of *FveTRM5* in promoting organ elongation. We showed that *FveTRM5* regulates fruit shape by influencing both cell elongation and cell division. Our findings provide an excellent target gene for breeding new fruit shape cultivars and exploring fruit shape regulatory mechanisms in strawberry.

## 2. Materials and methods

### 2.1 Plant materials and growth conditions

Three *F. vesca* accessions, Yellow Wonder (white-fruited), Ruegen (red-fruited) and Hawaii 4 (H4, white-fruited), were used as the WT in this study. All plants were grown in a growth room under a light intensity of 100 μmol m^−2^ s^−1^ with a photoperiod of 16 h light and 8 h dark at 22 °C. For EMS mutagenesis, 0.4% EMS (Sigma-Aldrich; cat no. M0880) was used for seed treatment for 8 h at room temperature with gentle shaking. Mutants were screened in the M_2_ generation.

### 2.2 Phenotypic analysis

Flowers or fruits from three to five plants of each genotype were analyzed. The maximum length and width of each tissue were measured using Image J software, and the shape index was calculated as the ratio of length to width. Three fruits at 0 DAP of Ruegen, H4, *rf* and *FveTRM5*-OE were used for paraffin sectioning. The samples were quickly placed in a mixed fixative of formalin-acetic acid-ethanol (FAA) (70 % ethanol, 5 % formaldehyde and 5 % acetic acid). Paraffin embedding and sectioning were performed as previously described [34]. The section thickness was 10 μm for each fruit. For cell shape analysis, 20 cells in the pith with clear shapes were selected from one section of each fruit, resulting in 60 cells for each genotype. For cell number analysis, the cells in the widest part of the receptacle were counted over the entire receptacle from two sections of each fruit, resulting in 6 samples for each genotype.

### 2.3 Gene isolation of the *rf* mutant

The *rf* mutant was crossed with wild-type Ruegen to generate an F_2_ population. Equal amounts of young leaves from 11 F_2_ mutant and 20 F_2_ wild type plants were pooled. Genomic DNA was extracted using a CTAB method. Genome sequencing was performed on the Illumina HiSeq X Ten platform (Novogene, Beijing) and analyzed as previously described [35]. DNA-seq reads from the two pools were aligned to the woodland strawberry reference genome ver. 4 using BWA-MEM with default parameters [36, 37]. The SAM file was further transformed into a sorted bam file and duplicate reads were removed using SAMtools [38]. The SNPs in *F. vesca* were called using GATK HaplotypeCaller [39] with ‘--min-base-quality-score 20; --minimum-mapping-quality 20’ and then filtered with the following parameters: QD < 2.0 || FS > 60.0 || MQ < 40.0 || MQRankSum < -12.5 || ReadPosRankSum < -8.0 || -cluster 2 -window 20. Only G-A or C-T sites were retained for further analysis. The ΔSNP index was first calculated by subtracting the SNP frequency between two pools, and then a sliding window analysis with a 300-kb window and 100-kb step was performed to calculate the average ΔSNP index. SNPs located in the CDS region were extracted using BEDTools [40]. The candidate mutation was examined in individual F_2_ mutants by PCR amplification and Sanger sequencing.

### 2.4 Phylogenetic analysis

The FveTRM proteins in woodland strawberry were identified from the PLAZA website (https://bioinformatics.psb.ugent.be/plaza/versions/plaza_v5_dicots/) grouped together with AtTRMs. Protein sequences were obtained from TAIR for Arabidopsis (Arabidopsis.org), Sol Genomics Network for tomato (solgenomics.net), and Cucurbit Genomics Data for cucumber (cucurbit genomics.org). An unrooted phylogenetic tree was constructed using MEGA7 with the neighbor-joining statistical method and bootstrap analysis (1000 replicates).

### 2.5 Plasmid construction

Genomic DNA or total RNA was extracted from fruits or young leaves of Ruegen and used for gene amplification. For overexpression in woodland strawberry, the full-length coding sequence of *FveTRM5* was cloned into pENTR1A and inserted into the binary vector pK7WG2D. For overexpression in Arabidopsis, the full-length coding sequence of *FveTRM5* was inserted into pRI101 at the *Sal*I and *BamH*I site and fused with GFP using the ClonExpress II One Step Cloning Kit (Vazyme, C112-01). For *FveTRM5pro*:*FveTRM5*, 304 bp upstream of the translation start site, 3,219 bp of the gene body (exons and introns), and 999 bp downstream of the stop codon of *FveTRM5* were inserted into the binary vector pCAMBIA1300 at the *Sal*I and *Bam*HI sites. For subcellular localization, the coding sequence of *FveTRM5* was cloned into the binary vector pH7LIC5.0 fused to N-terminal GFP at the *Sal*I and *Stu*I sites using the ClonExpress II One Step Cloning Kit (Vazyme, C112-01). Primers are listed in Table S1.

### 2.6 Stable transformation in woodland strawberry

Transformation in woodland strawberry was performed as previously described [41]. The overexpression and complementation constructs were transformed separately into woodland strawberry variety H4. During transformation, positive overexpression transgenic calli and regenerated plants were selected using 10 mg L^−1^ kanamycin and GFP fluorescence as examined under a fluorescence dissecting microscope (Microshot Technology Ltd, Guangzhou, China, MZX81). Positive transgenic calli of *FveTRM5*pro:*FveTRM5* were selected on the medium containing 4 mg L^−1^ hygromycin.

### 2.7 Stable transformation in Arabidopsis

Arabidopsis Col-0 was transformed with *Agrobacterium tumefaciens* GV3101 using the floral-dip method. The transgenic lines were selected on half strength MS (M5524, Sigma) with 100 mg L^-1^ kanamycin in the T_1_ generation.

### 2.8 Subcellular localization analysis

*Nicotiana benthamiana* leaves were infiltrated on the abaxial side with the *A. tumefaciens* strain GV3101 cells containing the construct and the silencing suppressor p19 in a 1:1 ratio. The *35S*:*RFP-AtTUA5* construct was co-infiltrated as a microtubule marker [42]. Agrobacterium cells were harvested and resuspended in the buffer (10 mM MgCl_2_, 10 mM MES, pH 5.6 and 150 mM acetosyringone at pH 5.6) and adjusted to an OD_600_ of 0.6. The solutions were incubated for 2 hours at room temperature without shaking before infiltration. Fluorescence signals were acquired four days after infiltration using a Leica TCS SP8 inverted microscope. Excitation wavelengths were 488 nm for GFP and 552 nm for RFP, and emission was detected at 505-550 nm for GFP and 590-640 nm for RFP.

### 2.9 Quantitative RT-PCR

Total RNA was extracted by using a HiPure Plant RNA Mini Kit (Magen, Guangzhou, China; cat no. R4151) and reverse transcribed to cDNA using HiScript III RT SuperMix for qPCR (+gDNA/wispr, Vazyme, Najing, China; cat no. R323). qPCR was performed using a Quant Studio 7 Flex system (Applied Biosystems, Waltham, MA, USA). The expression level of each gene was calculated using the 2^−ΔΔCT^ method. FvH4_1g05910 was used as an internal control. Three biological replicates were used for each sample, and three technical replicates were analyzed for each biological replicate. Primers are listed in Table S1.

### 2.10 RNA *in situ* hybridization

Shoot tips and flower buds at different developmental stages of wild-type H4 were collected and fixed in the 50% FAA fixative solution at 4 °C. A 200 bp fragment of the *FveTRM5* gene (215 to 414 bp in the coding sequence) was inserted into the pGEM-T vector at the *Nco*I and *Sal*I sites. The DIG RNA Labeling Kit (SP6/T7; Roche, Cat#11175025910) was used to synthesize DIG-labeled RNA probes by in vitro transcription. The sense probe was synthesized by T7 polymerase, and the antisense probe was synthesized by SP6 polymerase. The hybridization signals were detected using the DIG Nucleic Acid Detection Kit (Roche, Cat#11175041910). Slides were developed for 18 h in a dark moist container and stored in 1× TE after the reaction was stopped. Images were captured using a Leica microscope (DM6B) with a 10× or 20× optical adapter. Primers are listed in Table S1.

### 2.11 Statistical analyses

Statistical analyses were performed using the GraphPad Prism 8 software. Pairwise comparisons were made using Student’s *t*-test (ns, not significant; *, *P* < 0.05; **, *P* < 0.01).

## 3. Results

### 3.1 The *rf* mutant produces round or flat fruits in woodland strawberry

To identify genes regulating strawberry fruit shape, we found a *round fruit* (*rf*) mutant from the ethyl methanesulfonate (EMS)-mutagenized population of the woodland strawberry variety Yellow Wonder (YW, white-fruited). The *rf* mutant with red fruits was generated after crossing with the red-fruited variety Ruegen (Fig. S1A) [43]. The wild-type fruits were conical, whereas the *rf* fruits were nearly round or flat (Fig. 1A). Fruit measurements showed a significant reduction in fruit length and a significant increase in fruit width, resulting in a significantly reduced fruit shape index (length/width) in *rf* compared to wild type, but no difference in fruit weight (Fig. 1B). Other *rf* organs, such as petals, sepals, receptacles, stamens, and leaves, were also shorter or rounder than the wild type (Fig. 1C, D; Fig. S1B, C). Further observation showed that the *rf* receptacle was always shorter than the wild type from anthesis to fruit ripening (Fig. 1E). These results indicate that the *RF* gene plays an important role in regulating the shape of both fruit and other organs in woodland strawberry.

**Figure 1.**
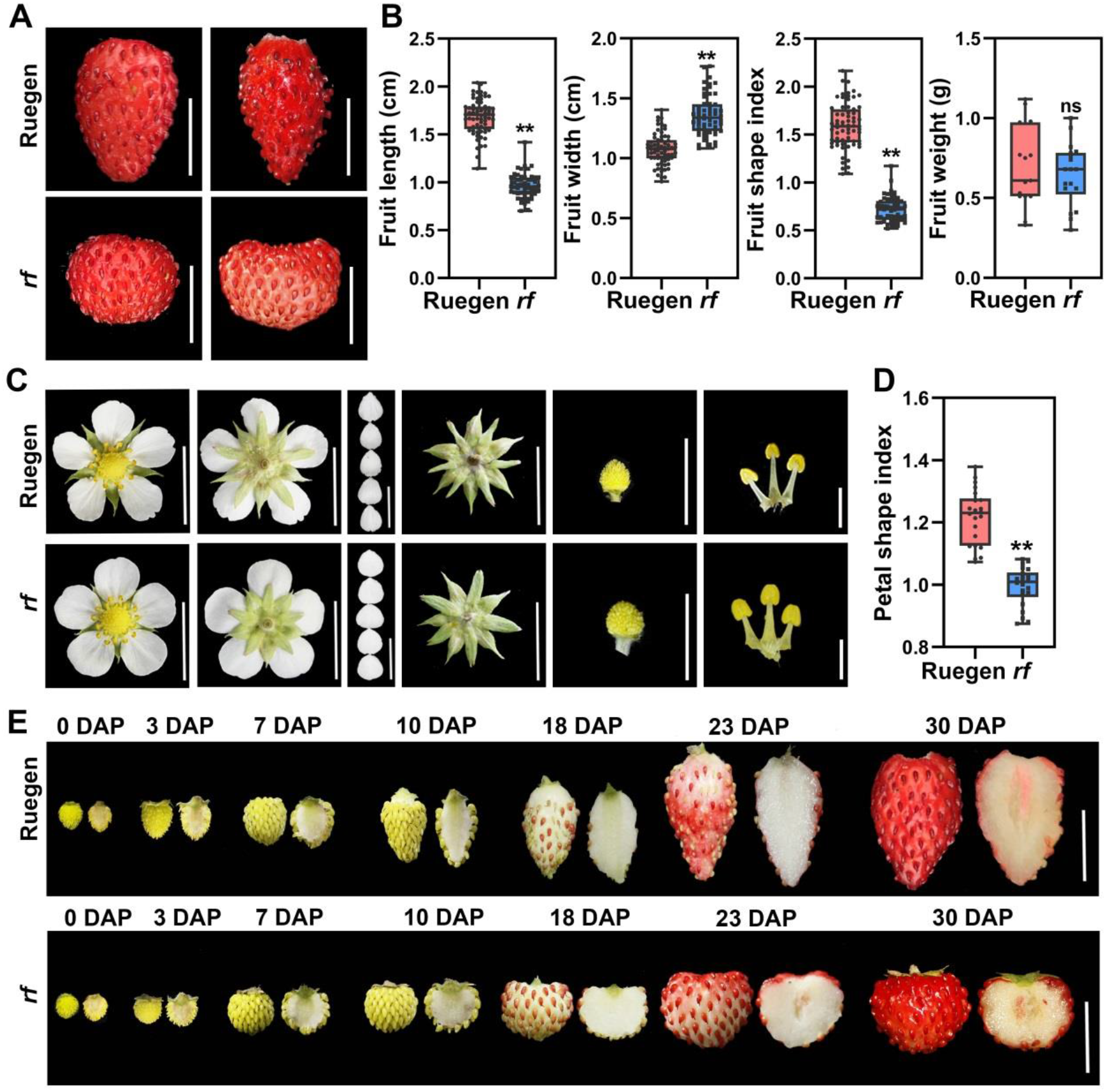
Fruit morphologies in round fruit mutants in in woodland strawberry. (A) Mature fruits of wild-type Ruegen and the *rf* mutant. Scale bars: 1 cm. (B) Fruit width, length, shape index and weight of wild-type Ruegen and *rf. n* > 20. (C) Images showing the floral tissues of wild-type Ruegen and the *rf* mutant. Scale bars: 2 mm for stamens and 1 cm for others. (D) Petal shape index of wild-type Ruegen and the *rf* mutant. *n* > 20. (E) Fruits of wild-type Ruegen and *rf* at different developmental stages from anthesis to ripening. DAP, day after pollination. Scale bars: 1 cm. For statistical analysis, data are the mean ±SD; ^**^, *P* < 0.01; ns, not significant, Student’s *t*-test.

### 3.2 *FveTRM5* is the causal gene of the *rf* mutant

To identify the causal gene, the *rf* mutant was crossed with the wild-type Ruegen, resulting in F_1_ progeny that all exhibited the wild-type phenotype. These F_1_ plants were then self-pollinated to produce a segregating F_2_ population. In this F_2_ population, 81 plants produced conical fruits, and 27 plants produced round fruits, suggesting a monogenic recessive mutation responsible for *rf*. Bulked genome resequencing and data analysis identified a highly linked peak on chromosome 2 (Fig. 2A). The following criteria were used to filter the SNPs: (1) G-to-A or C-to-T transition; (2) with an index of 100% in the mutant library, < 50% in the WT library; (3) located in the coding sequence and causing nonsynonymous mutations in the protein [35]. After filtering, seven candidates were identified in the region from 16.48 to 20.03 Mb (Table S2). One of the candidates is the C-to-T mutation in the second exon of FvH4_2g22810 (annotation ver. 4), which causes a premature stop codon at amino acid residue 266 (Fig. 2B). This was further confirmed to be homozygous in 52 F_2_ mutants by Sanger sequencing, in which the mutation in FvH4_2g22810 co-segregated with the round fruit phenotype. The protein sequence of FvH4_2g22810 shared a high similarity with Arabidopsis TON1 RECRUITING MOTIF 5 (AtTRM5), tomato SlTRM5, and cucumber CsTRM5 (Fig. 2C) [8, 14, 15]. Therefore, this gene was named *FveTRM5*. We found a total of 18 TRM family members in the woodland strawberry genome with distinct expression patterns (Fig. S2A, B). The AtTRM1-5 subclade contains three woodland strawberry genes, including *FveTRM1* (FvH4_6g46710), *FveTRM4* (FvH4_5g13690) and *FveTRM5* (Fig. 2C). When examined by RT-qPCR, *FveTRM5* was significantly downregulated in the *rf* fruits at 7 DAP (days after pollination) compared to wild type (Fig. 2D), suggesting nonsense-mediated RNA decay.

**Figure 2.**
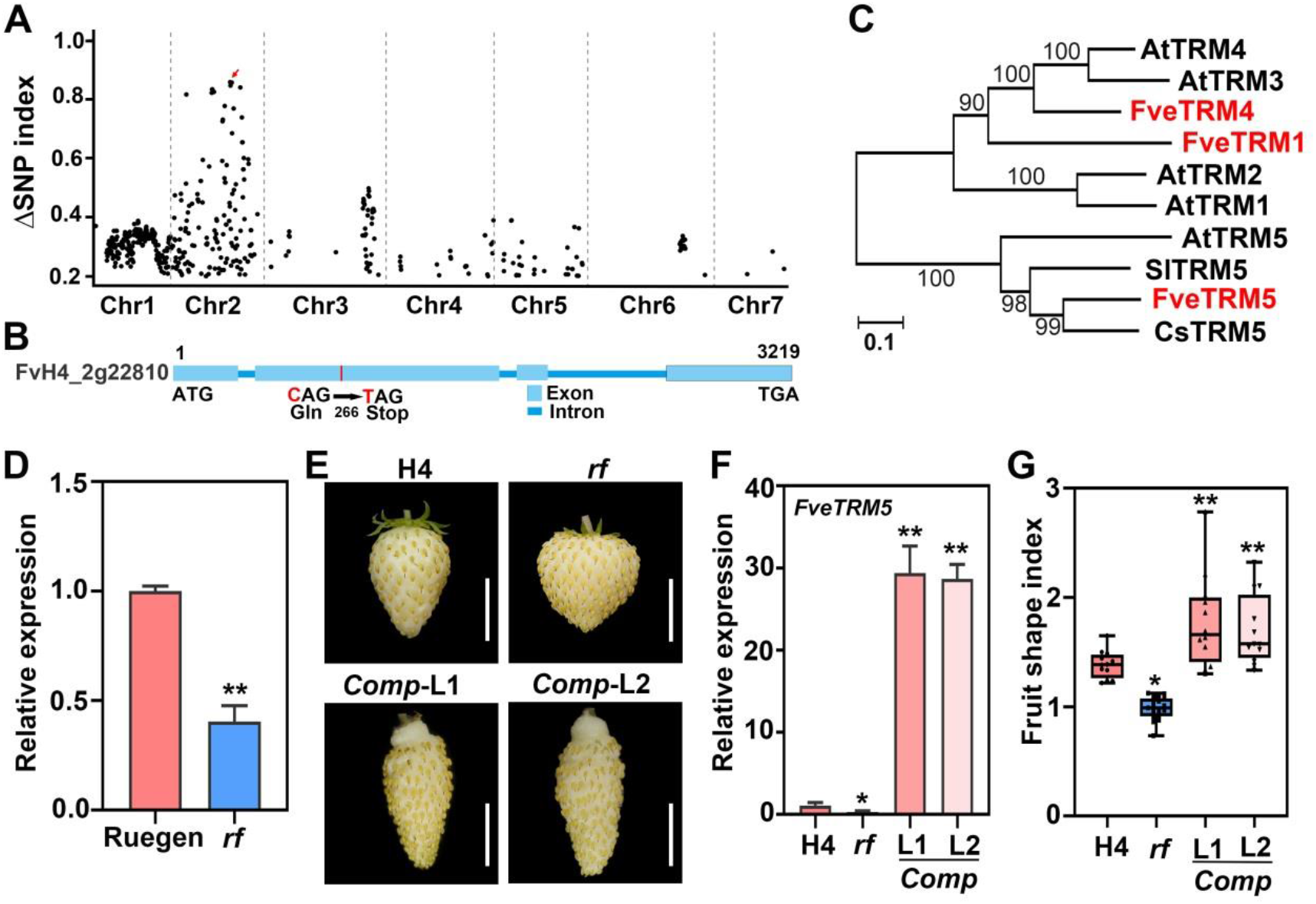
Isolation and characterization of *FveTRM5* in woodland strawberry. (A) Diagram showing the SNPs associated with *rf* along the chromosomes. The X axis represents seven chromosomes in woodland strawberry. The Y axis indicates the differences in allele frequencies between the wild type and *rf* mutant pools. The arrow indicates the linked peak on chromosome 2. (B) Diagram showing the candidate causal mutation in FvH4_2g22810 for *rf*. The red letters indicate the candidate point mutation. (C) Phylogenetic tree of the TRM proteins in woodland strawberry (Fve), Arabidopsis (At), tomato (Sl), and cucumber (Cs) in the AtTRM1-5 subclade using full-length sequences. Bootstrap values at the nodes are percentages of 1,000 replicates. The branch length indicates the number of substitutions per site. (D) Relative expression level of *FveTRM5* in wild-type Ruegen and *rf* fruits (7 DAP) examined by RT-qPCR. *n* = 3. (E) Images showing the mature fruits of wild-type H4, *rf*, and two complementation lines (*Comp*-L1 and *Comp*-L2) in woodland strawberry. Scale bars: 1 cm. (F) Relative expression level of *FveTRM5* in wild-type H4, *rf, Comp*-L1 and *Comp*-L2 leaves examined by RT-qPCR. *n* = 3. (G) Fruit shape index of H4, *rf, Comp*-L1 and *Comp*-L2. *n* > 10. For statistical analysis, data are the mean ±SD; ^*^, *P* < 0.05; ^**^, *P* < 0.01; Student’s *t*-test.

To genetically validate the gene cloning result, the complementation construct *FveTRM5pro*:*FveTRM5* was generated and stably transformed into the wild-type Hawaii 4 (H4) for higher transformation efficiency. We obtained a total of 28 transgenic lines with either wild-type or elongated fruits in the T_0_ generation. We selected 2 lines (L1 and L2) with elongated fruits to cross with the *rf* mutant. In the F_2_ generation, the transgenic plants with the homozygous *rf* mutation were identified and named *Comp*-L1 and *Comp*-L2 (Fig. 2E). RT-qPCR showed that the expression level of *FveTRM5* in *Comp*-L1 and *Comp*-L2 was 28 to 30 times higher than in the wild type (Fig. 2F). The fruit shape index of *Comp*-L1 and *Comp*-L2 was significantly higher than that of *rf* and wild type (Fig. 2G), indicating that *FveTRM5* could rescue the fruit shape defect in *rf*. Taken together, these results indicate that *FveTRM5* is the causal gene of the *rf* mutant.

### 3.3 Overexpression of *FveTRM5* results in elongated organs in Arabidopsis and woodland strawberry

To determine the function of *FveTRM5, FveTRM*5 driven by the 35S promoter was first transformed into WT Arabidopsis. Twenty-five independent *FveTRM5*-OE lines were obtained in the T_1_ generation with much narrower and longer leaves (Fig. S3A). *FveTRM5* expression was confirmed in two transgenic Arabidopsis lines (Fig. S3B). Compared to the wild type, overexpression of *FveTRM5* resulted in sufficiently elongated organs, including petals, stamens, gynoecia, siliques, and seeds (Fig. S3C, D). These results suggest that *FveTRM5* could promote organ elongation in alternative species.

To further clarify the function of *FveTRM5* in strawberry, the *FveTRM5*-overexpression construct was also stably transformed into the wild-type woodland strawberry strain H4. We obtained 10 independent *FveTRM5* overexpressing lines. Two lines (L2 and L3) were carefully characterized at T_0_. Both lines produced much thinner mature fruits and floral organs, such as petals and receptacles (Fig. 3A). RT-qPCR confirmed that *FveTRM5* was expressed at 177-fold and 100-fold higher levels in these two lines than in the wild type (Fig. 3B). Fruit shape measurements showed that fruit length was significantly greater and fruit width was significantly smaller in *FveTRM5*-OE than in the wild type, resulting in a higher fruit shape index (Fig. 3C). It should be noted that the fruit weight also decreased significantly in *FveTRM5*-OE, mainly due to the extreme reduction in fruit width. The change in fruit shape in *FveTRM5*-OE started at anthesis and remained the same until ripening (Fig. 3D). *FveTRM5*-OE also showed an increase in the petal shape index (length/width) (Fig. 3E). In addition, *FveTRM5* overexpression resulted in narrower leaves (Fig. S4A-C). Taken together, *FveTRM5* is an important regulator of organ shape in woodland strawberry and may exert similar phenotypes in other plants.

**Figure 3.**
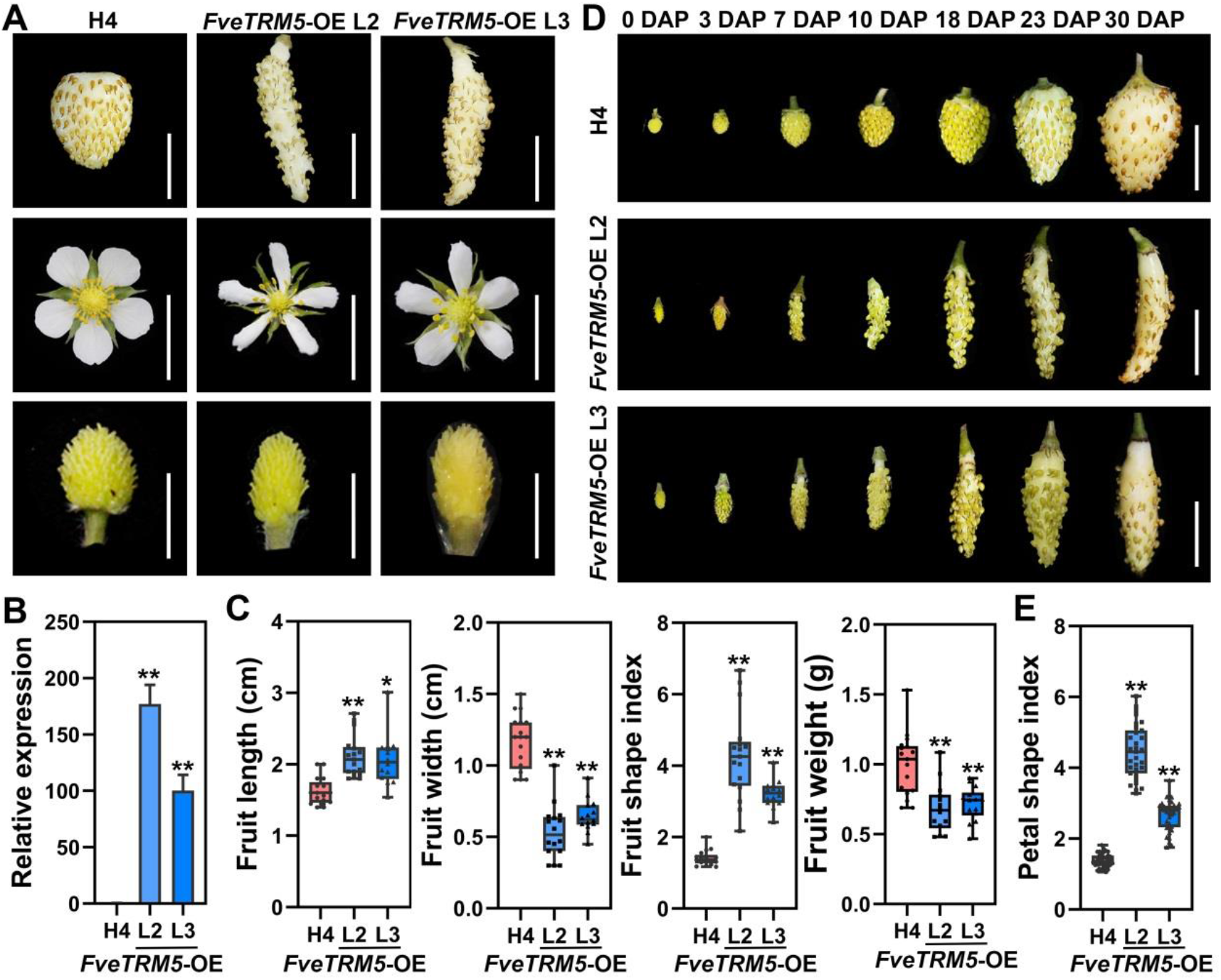
Phenotypic characterization of the *FveTRM5-OE* transgenic lines in woodland strawberry. (A) Images showing the fruits and floral organs of two *FveTRM5*-OE transgenic lines (L2 and L3) in woodland strawberry. Scale bars: 1 cm. (B) Relative expression levels of *FveTRM5* in the leaves of wild-type H4 and *FveTRM5*-OE (L2 and L3) examined by RT-qPCR. *n* = 3. (C) Fruit length, width, shape index and weight of H4 and *FveTRM5*-OE (L2 and L3). *n* > 15. (D) Fruits of wild type H4 and two independent *FveTRM5*-OE lines (L2 and L3) at different developmental stages from anthesis to ripening. DAP, days after pollination. Scale bars: 1 cm. (E) Petal shape index of H4 and *FveTRM5*-OE (L2 and L3). *n* > 20. For statistical analysis, data are the mean ±SD; ^*^, *P* < 0.05; ^**^, *P* < 0.01, Student’s *t*-test.

### 3.4 *FveTRM5* is highly expressed in developing organs

Based on the woodland strawberry transcriptome database [44], *FveTRM5* was widely expressed in flower and fruit tissues, such as flower meristems (FM), carpels, embryos, and receptacles (cortex and pith), as well as shoot apical meristems (SAM), leaves and roots (Fig. 4A). When examined by RT-qPCR, *FveTRM5* expression was indeed higher in leaves, flower buds and young fruits, moderate in roots, but strongly decreased in mature fruits at 30 DAP (Fig. 4B). We further examined the expression pattern of *FveTRM5* in shoot tips and floral buds by RNA *in situ* hybridization. Flower stages were designated as previously described [45]. Strong signals of *FveTRM5* were detected in the SAM, leaf primordia, FM, receptacle meristem and the four whorls of floral organs at flower stages 1-13, no signal was detected by hybridization with the *FveTRM5* sense probe (Fig. 4C).

**Figure 4.**
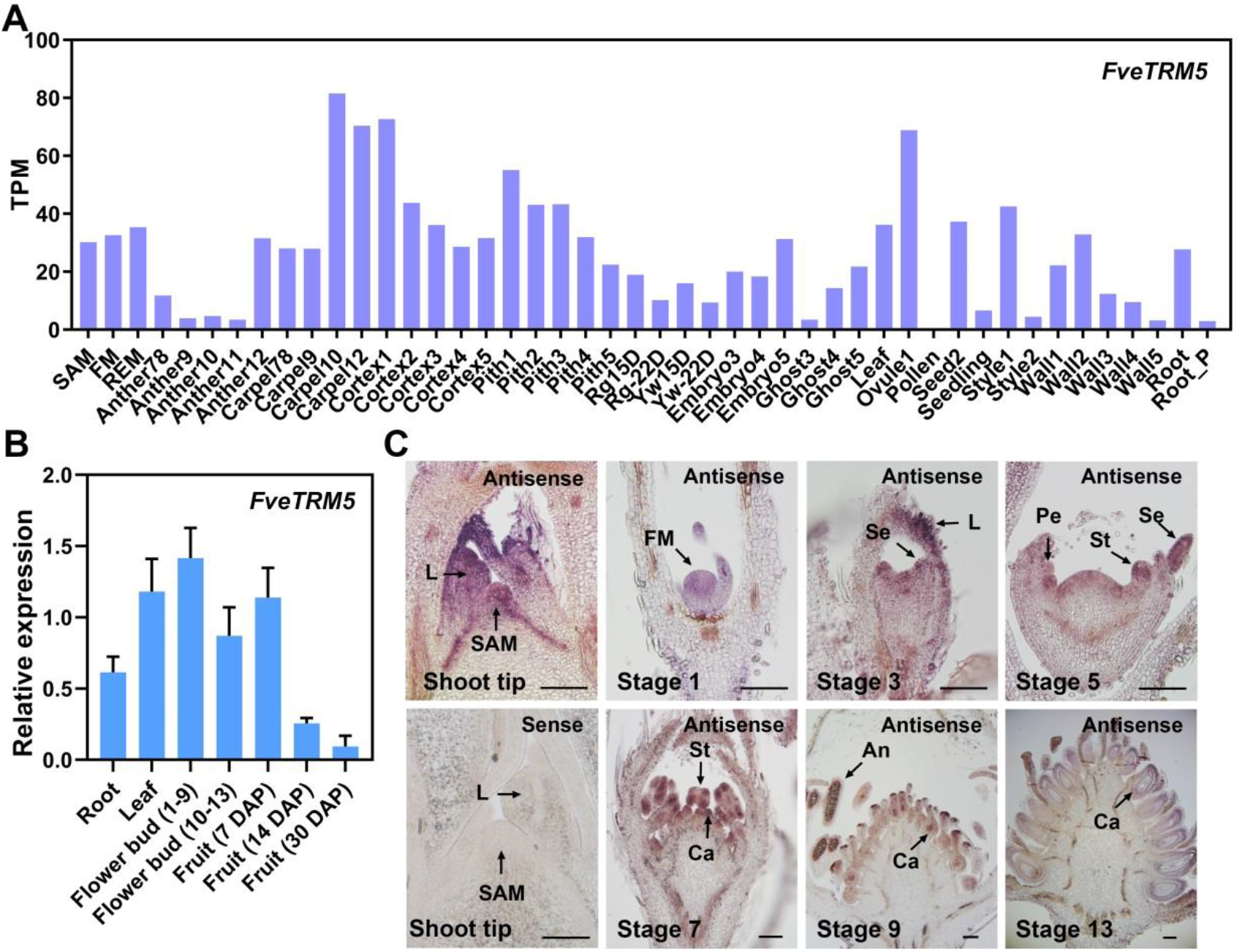
Expression pattern of *FveTRM5* in woodland strawberry. (A) Expression pattern of *FveTRM5* in woodland strawberry as indicated by transcripts per million (TPM) values obtained from the transcriptome database (Li et al., 2019). (B) Relative expression levels of *FveTRM5* in the tissues of wild-type Ruegen examined by RT-qPCR. *n* = 3. DAP, days after pollination. (C) Expression pattern of *FveTRM5* in longitudinal sections of wild-type shoot tips and flower buds at different stages by RNA *in situ* hybridization. SAM, shoot apical meristem; L, leaf; FM, floral meristem; Se, sepal; St, stamen; Pe, petal; Ca, carpel; An, anther. Scale bars: 100 μm.

### 3.5 *FveTRM5* regulates cell expansion and cell division in strawberry fruit

As the change in fruit shape remains the same throughout development, the cell morphology in the receptacles of *rf, FveTRM5*-OE and the wild types (Ruegen and H4) at anthesis was examined by paraffin sectioning. Initial observations showed that the receptacle cells appeared shorter and smaller in *rf* and longer in *FveTRM5*-OE (Fig. 5A). Cell measurements showed that *rf* receptacle cells were significantly shorter than wild type, whereas *FveTRM5*-OE receptacle cells were significantly longer than wild type (Fig. 5B), suggesting that FveTRM5 positively regulates cell elongation in the longitudinal direction. In contrast, cell width was significantly decreased in *rf* and the two *FveTRM5*-OE transgenic lines, resulting in the significant change of cell shape index. In addition, there were more cell layers in *rf* and fewer cell layers in *FveTRM5*-OE in the transverse direction, suggesting that *FveTRM5* could inhibit cell division in the medial-lateral direction of the receptacle (Fig. 5B).

**Figure 5.**
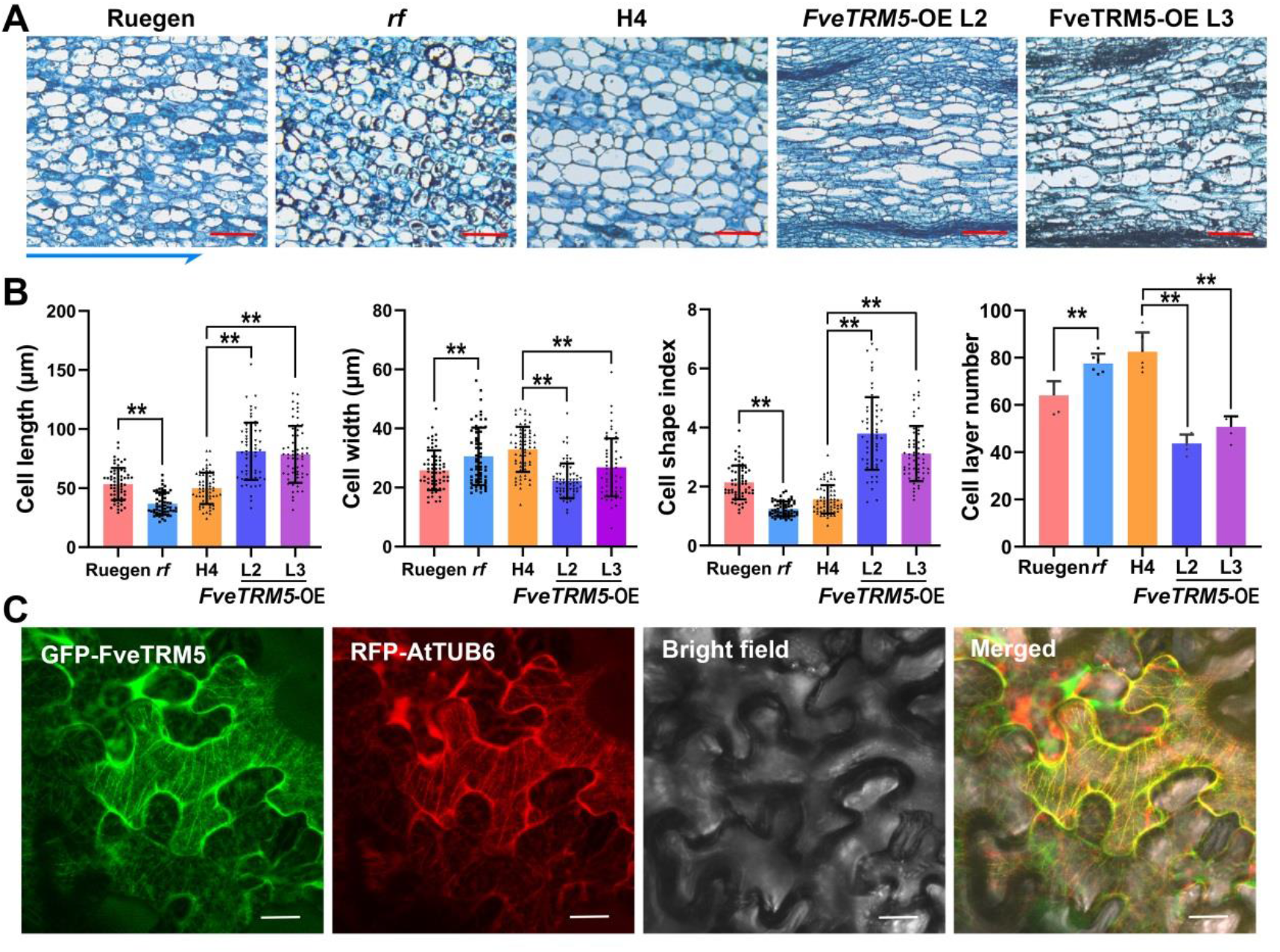
Changes in cell morphology in different materials and subcellular localization of FveTRM5. (A) Images showing cell shapes in the receptacle of wild-type Ruegen and H4, *rf*, and *FveTRM5-OE* (L2 and L3) at anthesis. The blue arrow indicates the longitudinal direction of the fruit. Scale bars: 100 μm. (B) Cell shape and number of cell layers in the transverse direction at the widest part of wild-type Ruegen and H4, *rf*, and *FveTRM5-OE* (L2 and L3) receptacles. For statistical analysis, data are the mean ±SD; ^**^, *P* < 0.01; student’s *t*-test. *n* = 60 for cell shapes and 6 for cell layers. (C) Subcellular localization of GFP-FveTRM5 in *Nicotiana benthamiana* leaves. Green indicates GFP fluorescence, and red indicates RFP fluorescence of the microtubule marker RFP-AtTUB6. Scale bars: 10 μm.

Previous studies have shown that some TRM proteins co-localize with microtubules, a major component of the cytoskeleton [14]. To test this possibility, GFP-FveTRM5 and the microtubule marker RFP-AtTUB6 were simultaneously infiltrated into the *Nicotiana benthamiana* leaves for transient expression. The RFP-AtTUB6 signals were shown as lines, indicating the localization of microtubules (Fig. 5C). GFP-FveTRM5 signals colocalized with RFP-AtTUB6 on microtubules (Fig. 5C). This localization is similar to that of SlTRM5, which has been implicated in fruit shape regulation [14, 46]. Taken together, these results indicate that *FveTRM5* regulates strawberry fruit shape by controlling both cell expansion and cell division.

## 4. Discussion

Cultivated strawberries produce fruits with a wide variety of shapes, but those with round fruits are relatively rare. In this study, we identified an EMS mutant called *round fruit* (*rf*) in woodland strawberry. Gene cloning and genetic analysis revealed that *FveTRM5* is the causal gene for this mutant. Overexpression of *FveTRM5* resulted in elongated organs in both woodland strawberry and Arabidopsis, suggesting a conserved role in different species. Furthermore, *FveTRM5* can inhibit transverse cell division and promote longitudinal cell elongation in the fruit receptacle. This work provides compelling evidence for the role of *FveTRM5* in fruit shape control and makes it a strong candidate for fruit shape manipulation in strawberry and other fruit crops.

The TRM proteins in the AtTRM1-5 subclade promote elongation of fruits as well as other organs in a wide range of plant species [9-11, 14, 15]. There are three members in this subclade in the woodland strawberry genome (Fig. 2C). In this study, we have demonstrated the important functions of *FveTRM5* in fruit shape regulation. It remains to be tested whether *FveTRM1* and *FveTRM4* have similar and redundant functions. This possibility is supported by the tomato mutants of *SlTRM3/4* and *SlTRM5*, which show additive effects in shortening the *ovate Slofp20* fruits [46]. In addition, mutations in *SlTRM19* and *SlTRM17/20a* from another subclade could enhance fruit elongation in the *ovate Slofp20* double mutant, suggesting that the TRM proteins from different subclades may also have redundant or opposing functions. This finding extends the range of potential TRMs involved in the fruit shape control.

The TRM proteins promote organ elongation by increasing cell division in the longitudinal direction and decreasing cell division in the transverse direction [9, 14, 15]. Our results showed that *FveTRM5* not only inhibited cell division in the transverse direction, but also decreased cell width and increased cell length, a result of anisotropic cell expansion (Fig. 5). Although TRM proteins can be localized on microtubules or in the cytosol [8], SlTRM5 is particularly localized on microtubules [14, 46]. FveTRM5 also localizes to microtubules, similar to SlTRM5. The TRM proteins often interact with OFPs to regulate organ shape [6, 14]. Thus, protein interaction assays may help to find the FveOFPs involved in the fruit shape regulation.

Fruit shape is finely regulated by plant hormones, such as auxin and gibberellic acid (GA), as well as other stimuli. In tomato, whole-plant application of exogenous auxin before anthesis results in elongated ovaries and fruits, with an increased number of cells along the longitudinal axis and enlarged cells in most tissues of the ovary [47]. GA_3_ treatment in tomato results in elongated fruits, whereas application of the GA inhibitor paclobutrazol results in more flattened fruits [48, 49]. In woodland strawberry, auxin application results in rounder fruits, whereas GA treatment leads to thinner fruits [30]. The hormone pathways are known to cross-talk with the *OFP* and *IQD* family genes [6, 47, 50, 51], which may indirectly interfere with TRM functions. In addition to hormones, mutation of the red photoreceptor FvePhyB results in fruits with higher shape index [52]. Some upstream regulators of the *TRM* genes have been reported. For example, the bHLH transcription factors Leaf-related Protein 1 (LP1) and LP2 can directly regulate the expression levels of *AtTRM1* and *AtTRM2* to induce longitudinal cell elongation in Arabidopsis [53]. OsSPL16 can directly bind to the *GW7* (ortholog of *AtTRM1*) promoter to inhibit its expression in rice [10]. Whether and how *FveTRM5* is regulated by hormones and light signaling remains to be investigated.

In conclusion, we have reported a key player in the regulation of strawberry fruit shape, FveTRM5, which influences cell division and cell elongation in fruit tissues. Based on the genetic studies, we believe that it is possible to knock out *FveTRM5* to produce round fruits. However, *FveTRM5* cannot be overexpressed to high levels as this will reduce the biomass. These findings can provide a theoretical basis for breeding new fruit shape cultivars in strawberry and enrich our knowledge of fruit shape control in fruit crops.

## Data Availability

All relevant data and vectors that support the findings of this study are available from the corresponding author, upon request.

## Author Contributions

CK conceived and designed the experiments; ZZ and LW performed the experiments; QG created the F_2_ population of *rf*; SH analyzed the sequencing data; ZZ and CK wrote the manuscript. All the authors have read and approved the manuscript.

## Acknowledgments

The authors thank Dr. Pengwei Wang (Huazhong Agricultural University) for providing the microtubule marker used for subcellular localization.

## Funding

This work was supported by the National Natural Science Foundation of China (32172539).

## Declaration of Competing Interest

Patent entitled “The *RF* gene regulating strawberry fruit shape and its application (ZL202210171639.0)” has been authorized in China. CK and LW are listed as inventors.

## Supplementary data

**Figure S1**. Leaf phenotypic characterization of the *rf* mutant in woodland strawberry.

**Figure S2**. Phylogenetic and expression pattern analysis of the *FveTRM* genes.

**Figure S3**. Phenotypic characterization of the *FveTRM5*-OE transgenic lines in *Arabidopsis*.

**Figure S4**. Leaf phenotypic characterization of the *FveTRM5*-OE transgenic lines in woodland strawberry.

**Table S1**. Summary of primers used in this study.

**Table S2**. The list of candidate SNPs in the EMS mutant *rf*.

