## Supplementary data for "*FveTRM5* plays a critical role in regulating fruit shape in woodland strawberry"

**Supplementary information**

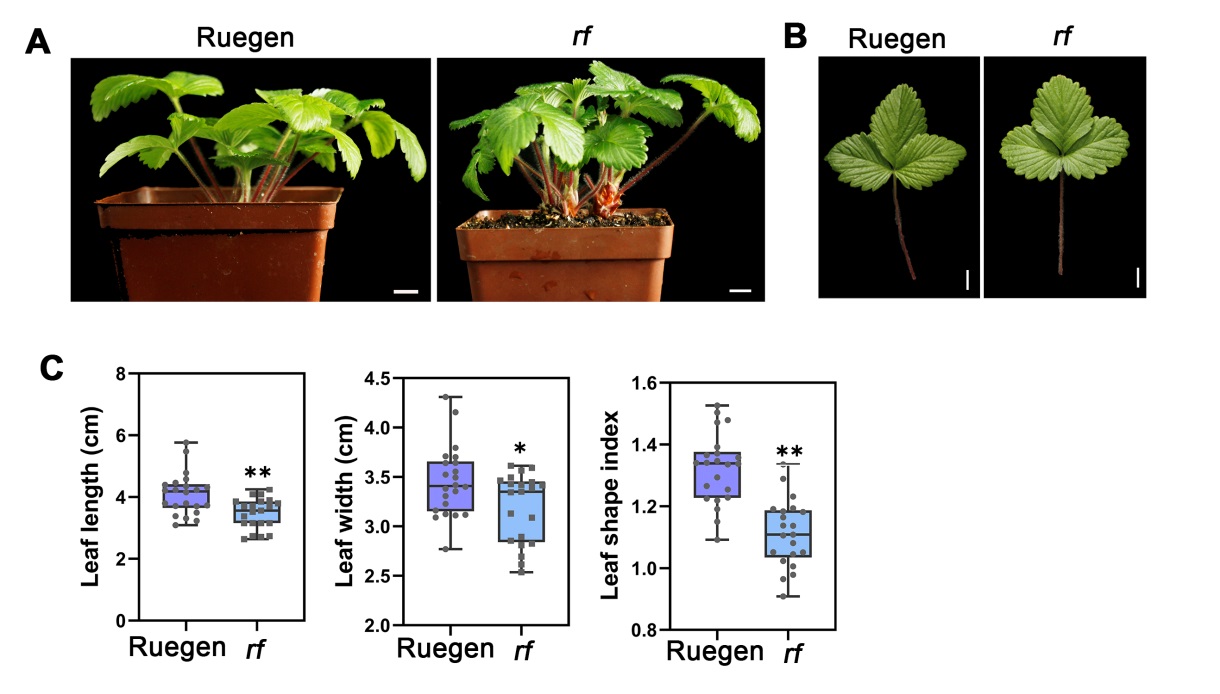

**Figure S1. Leaf phenotypic characterization of the *rf* mutant in woodland strawberry.**

(A) Images of wild type (Ruegen) and *rf* plants. Scale bars: 1 cm. (B) Adult leaves of Ruegen and *rf*. Scale bars: 1 cm. (C) Leaf length, width and shape index (length/width) of Ruegen and *rf*. *n* > 10. For statistical analysis, data are the mean *±*SD*;* *, P < 0.05; **, P < 0.01, Student’s t-test.

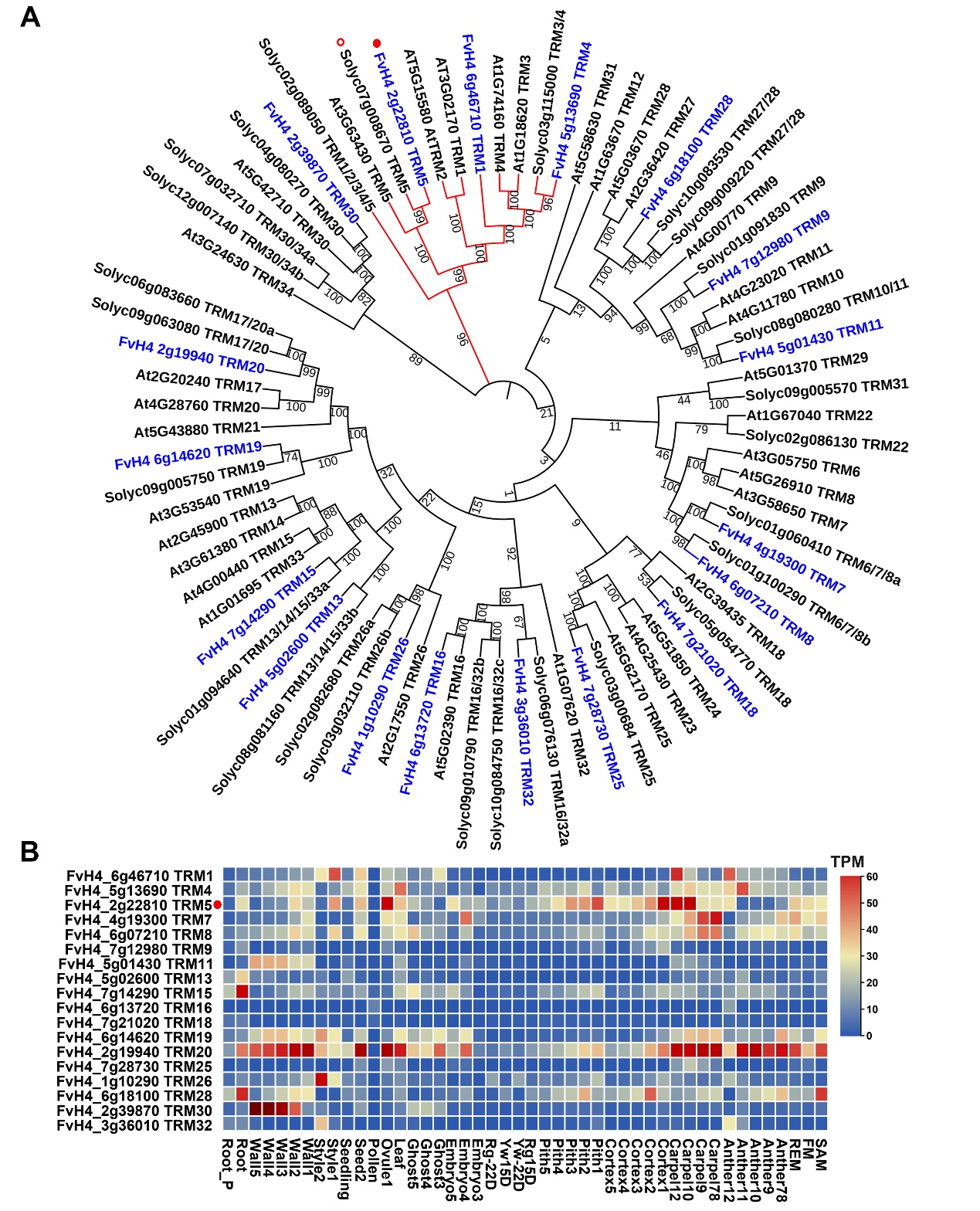

**Figure S2. Phylogenetic and** **expression pattern analysis of the *FveTRM* genes.**

(A) Phylogenetic tree of the TRM homologs from Arabidopsis (At), woodland strawberry (FvH4) and tomato (Solyc). Red branch indicates the AtTRM1-5 subclade, and blue labels indicate the FveTRM proteins. Red dot indicates FveTRM5, and red circle indicates SlTRM5. Bootstrap values at the nodes are percentages of 1,000 replicates. (B) Expression pattern of FveTRMs in woodland strawberry indicated by TPM (Transcript Per Million) values obtained from transcriptome database (Li et al., 2019). Red dot indicates FveTRM5.

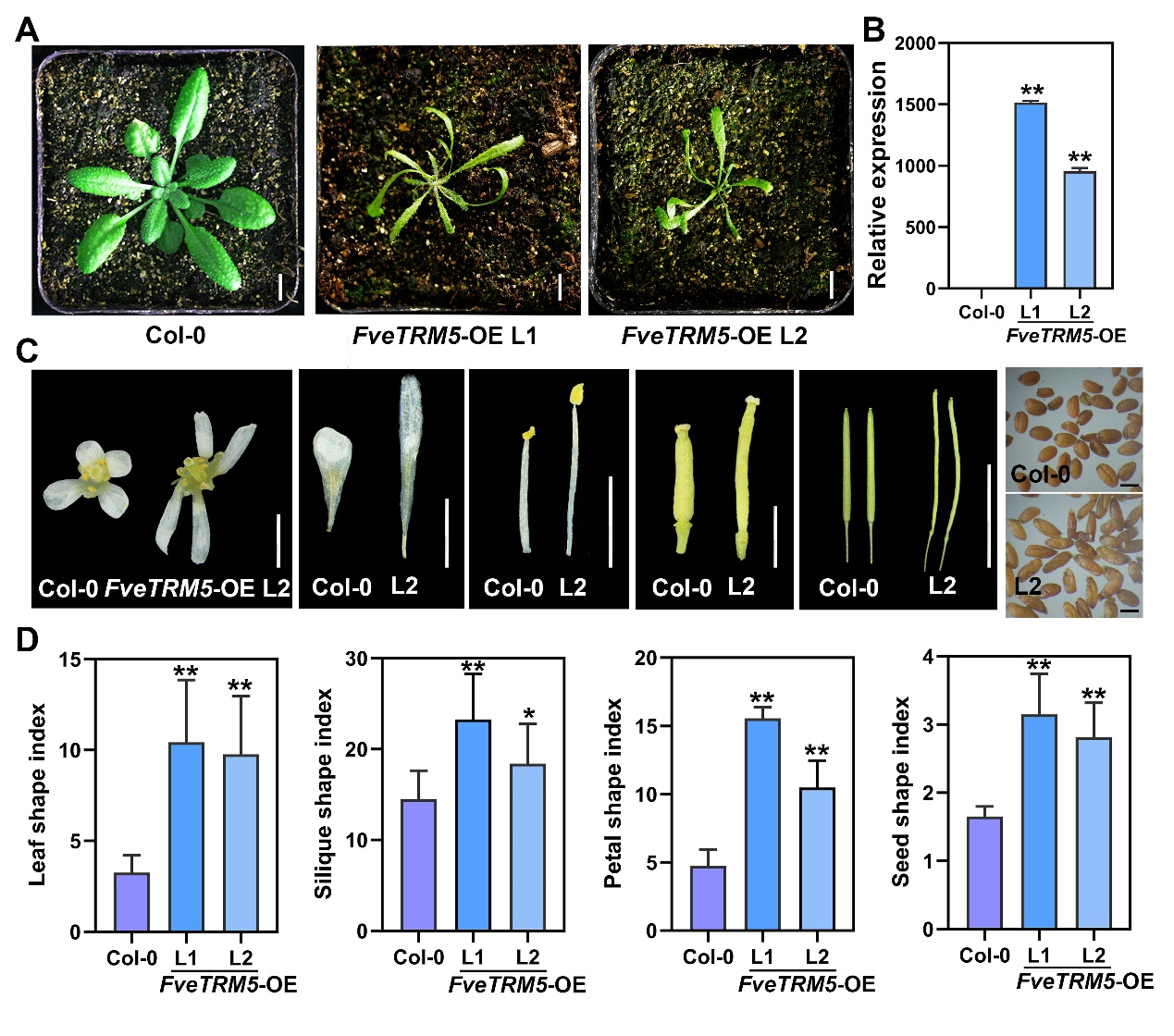

**Figure S3.** **Phenotypic characterization of the *FveTRM5-OE* transgenic lines in Arabidopsis.**

(A) Transgenic Arabidopsis plants overexpressing *FveTRM5* in the Col-0 background. Scale bars: 1 cm. (B) Relative expression levels of *FveTRM5* in the leaves of wild type and two independent transgenic lines in the T_1_ generation examined by RT-qPCR. (C) Floral organs and the seeds of the *FveTRM5-*OE transgenic Arabidopsis in the Col-0 background. From left to right, flowers, petals, stamens, carpels, siliques, and seeds are shown. Scale bars: 1 cm for siliques and 1 mm for others. (D) Bar plots showing the leaf shape index, silique shape index, petal shape index and seed shape index of wild type and the *FveTRM5-*OE lines in Arabidopsis. *n* > 10. For statistical analysis, data are the mean ±SD; *, *P* < 0.05; **, P < 0.01, Student’s t-test.

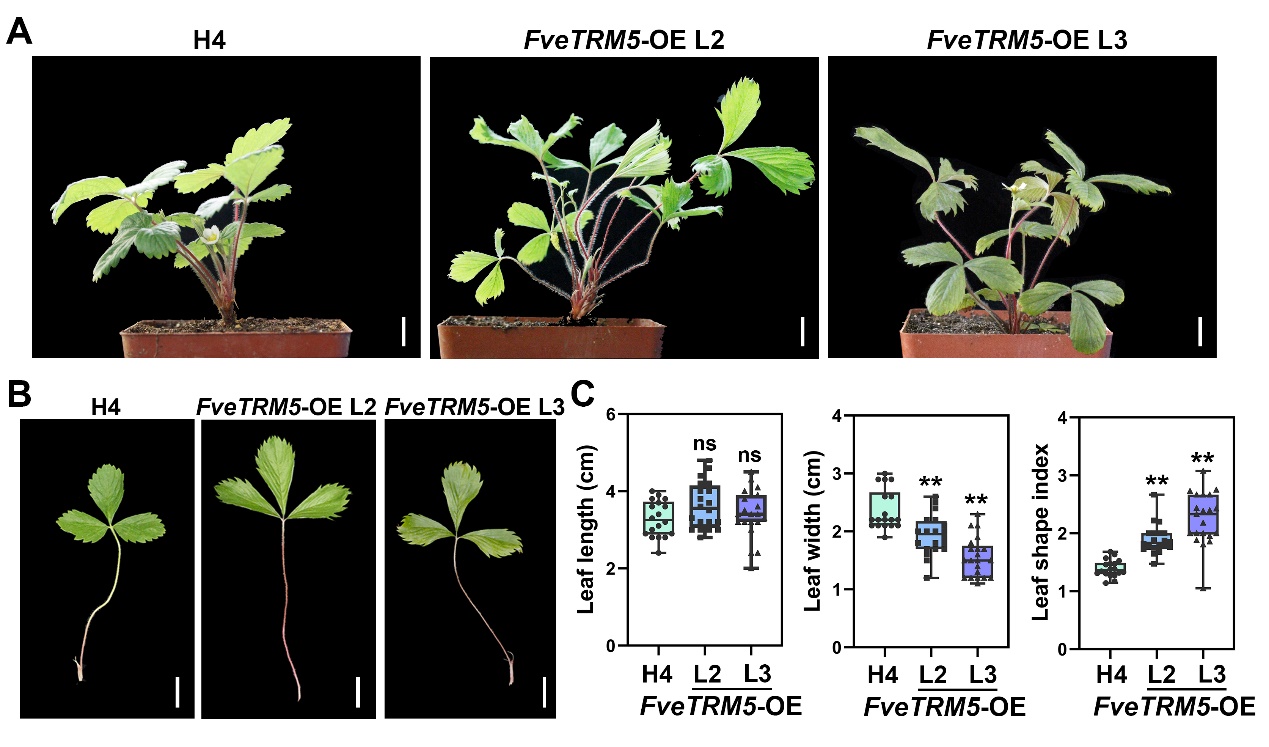

**Figure S4. Leaf phenotypic characterization of the *FveTRM5-OE* transgenic lines in woodland strawberry.**

(A) Images of wild type (H4) and two independent *FveTRM5*-OE transgenic (L2 and L3) plants. Scale bars: 1 cm. (B) Leaves of H4 and two independent *FveTRM5-OE* transgenic lines (L2 and L3) in the T_0_ generation. Scale bars: 1 cm. (C) Leaf length, width, and shape index (length/width) of H4 and two independent *FveTRM5-OE* transgenic lines (L2 and L3). *n* > 10. For statistical analysis, data are the mean *±*SD; **, P < 0.01; ns, not significant; Student’s t-test.

**Supplementary Table S1.** **Summary of primers used in this study.**

| **Primers for *rf* genotyping** | **Primer sequences (5’-3’)** |
| --- | --- |
| RF-F | CGAGGTCGTCGTGGAAGTTTTCGAG |
| RF-R | ACCTTTGAGCTGTAACGCTTCCAGG |
| **Primers for making the *FveTRM5* overexpression construct** | **Primer sequences (5’-3’)** |
| FveTRM5-pEN-F | AAAGGAACCAATTCAGTCGACATGACGACTGGGATGGTTCA |
| FveTRM5-pEN-R | TGGAAAAGGGAATTCGGTACCGAAGACCAACTTCCTACAAAGCGC |
| **Primers for making the *FveTRM5* complementation vector** |  |
| RF-com-F | TACGAATTCGAGCTCGGTACCTTCGTAAAAAGGAAGGGGCA |
| RF-com-R | CTTGCATGCCTGCAGGTCGACATACTCTGCTGGGGCATCAGAA |
| **Primers for the complementation material genotyping** |  |
| Com1-F | TTCAGGTCCGTGCACTCTTC |
| Com1-F | CTCTGCACCATTCCCCTGTT |
| Com2-F | CAGGAAACAGCTATGA |
| Com2-R | GTAAAACGACGGCCAGT |
| **Primers for making the construct for subcellular localization analysis** | **Primer sequences (5’-3’)** |
| FveTRM5-pH7-F | ATTACGCCGAGGTCATGACGACTGGGATGGTT |
| FveTRM5-pH7-R | TAGGGAAGAGGGAAGACCAACTTCCTACAAAGCG |
| **Primers for RT-qPCR** | **Primer sequences (5’-3’)** |
| FvH4_1g05910-F | AGCCTAACGCAGAGGTTCCAAA |
| FvH4_1g05910-R | GCAGCCCACATTGAAGGGTCTATAGT |
| FveTRM5-RT-F | GTTCACTCTCCCAAGGTTAGTT |
| FveTRM5-RT-R | CTCTGCTCTGTGGATGGTTATT |
| **Primers for RNA *in situ* hybridization** | **Primer sequences (5’-3’)** |
| FveTRM5-*Nco*I-F | GCATGCTCCCGGCCGCCATGGACCGCAAAACAAATCCCTGC |
| FveTRM5-*Sal*I-F | GAGCTCTCCCATATGGTCGACGCGGCATGTCTCTCGAAAAC |

**Supplementary Table S2.** **The list of candidate SNPs in the EMS mutant *rf*.**

| Chr | Position | DNA change | Amino acid change | %SNP in mutant library | %SNP in WT library | Gene ID (v2.2.a2) | Gene ID (v4.2.a2) |
| --- | --- | --- | --- | --- | --- | --- | --- |
| Fvb2 | 16482462 | CCC-CTC | pro-leu | G:0% A:100% | G:85% A:15% | gene17537 | FvH4_2g19380 |
| Fvb2 | 16590368 | GCT-GTT | ala-val | G:0% A:100% | G:81% A:19% | gene17557 | FvH4_2g19560 |
| Fvb2 | 17086023 | CTT-TTT | leu-phe | G:0% A:100% | G:74% A:26% | gene08354 | FvH4_2g20310 |
| Fvb2 | 17880038 | CTT-TTT | leu-phe | G:0% A:100% | G:75% A:25% | gene08083 | FvH4_2g21510 |
| Fvb2 | 18742176 | CAG-TAG | gln-stop | G:0% A:100% | G:89% A:11% | gene20276 | FvH4_2g22810 |
| Fvb2 | 18887568 | AGG-AAG | arg-lys | G:0% A:100% | G:82% A:18% | gene20346 | FvH4_2g23010 |
| Fvb2 | 20027554 | ACC-ATC | thr-lie | G:0% A:100% | G:69% A:31% | gene27776 | FvH4_2g24580 |
